# The effects of bacteriophage cocktail treatment on healthy gut microbiota: an *in vitro* human colon model study

**DOI:** 10.64898/2026.01.28.702202

**Authors:** Teagan L. Brown, Duncan Y.K. Ng, George M. Savva, Claire K.A. Elek, James A.D. Docherty, Ryan Cook, Rebecca Ansorge, Andrea Telatin, Elizabeth Kutter, Evelien M. Adriaenssens

## Abstract

The human gut microbiome is a complex community that plays an important role in health, where perturbations can result in dysbiosis and disease. Bacteriophages (phages) can provide treatment for bacterial gastrointestinal disease, and commercial preparations such as the Intesti bacteriophage cocktail can be taken orally to target bacterial pathogens. However, interactions between these phages and the native gut microbiota are understudied. To investigate the impact of phage treatment, we used simulated gut models seeded with healthy donor microbiota from three individuals, sequenced the DNA, and analysed the bacterial and viral portion from samples obtained over time. Each donor had a unique bacterial composition which diverged with time. When comparing phage treated to control samples, we observed that *Escherichia coli* abundance accounted for the largest portion of bacterial community variance and was more associated with the controls. The lower abundance in phage treated samples may have resulted from the lytic action of phages from the cocktail. Additionally, our analyses of the viral portion revealed a phage bloom exclusive to phage treated samples. A highly abundant phage in this bloom was matched with the Intesti bacteriophage cocktail, showed similarity to Enterobacteria phage phi92, and provided evidence of productive infection within the model. While we did observe fluctuations in relative abundance of additional viral sequences in the presence of the phage cocktail, these changes were often transient. Furthermore, we detected only slight differences to typical members of the virome, and low numbers of active prophages. Our experiments suggest that the phage cocktail had minimal interruption to the native gut microbiota within the model.

**Impact statement:** Bacteriophages are increasingly investigated and tested for their efficacy in treating infections and are a key component in fight against antimicrobial resistant bacterial infections. Because of their specificity, it has become almost a dogma to state that they do not alter the gut microbiome. We have now tested this in an *in vitro* study using a commercially available cocktail and real human faecal microbiota. We show minimal effects on the composition of the healthy microbiota with an individual-specific effect on *Escherichia coli* caused by productive infection of one phage in the cocktail.

## Introduction

The gut microbiome plays an essential role in human health and often is an indicator of disease development and progression^1^. Members of the phyla Bacteroidota and Bacillota dominate the bacterial portion of a healthy human adult gut, and studies report Bacteriodaceae, Clostridiaceae, Prevotellaceae, Ruminococcaceae, Bifidobacteriaceae, Lactobacillaceae and the Enterobacteriaceae as the most abundant families^2-4^. Enterotypes, and more recently enterosignatures, have been used to describe the bacterial community at the genus level, with the *Bacteroides* (western) and *Prevotella* (non-western) enterosignatures core to a healthy adult microbiome^5-7^.

The viral portion of the gut microbiome (virome), largely represented by bacteriophages (phages) is thought to play a significant role in the community by shaping the bacteria, and represents vast, often undescribed diversity^8, 9^. The development of improved viral detection tools^10-12^, human gut virome specific databases^9, 13-16^, and viral mining pipelines^17, 18^, have resulted in higher recovery of viruses from metagenomic datasets, and ongoing efforts continue to taxonomically classify the diversity^19^. A longitudinal study has shown that adult gut viromes are individualised and temporally stable, but are often dominated by dsDNA phages from the order *Crassvirales*, which show widespread distribution across multiple datasets^13, 20-22^, as well as *Microviridae* (ssDNA) phages^23^.

To investigate the dynamics of the gut microbiome in response to a range of stresses and conditions, *in vitro* model systems have been invaluable. These systems have been used to simulate bacterial dynamics in response to chemical and diet changes by static batch fermentation culture^24, 25^, continuous culture systems with mono or multi-stage compartments^26-28^, or the use of microfluidic organ-on-chip models^29, 30^. While acute changes to the gut microbiota composition can occur by several factors including the physical environment, nutrition, infection, medication and genetics, chronic or large shifts in the community can result in dysbiosis and disease states^31-33^. Diarrhoea, dysentery, enterocolitis and dysbacteriosis are common complaints of the gastrointestinal tract often associated with bacterial pathogens^34^. The use of antibiotics to treat diseases caused by these pathogens can have short-term (e.g. diarrhoea, *Clostridioides difficile* infection) but also profound and long-lasting effects on the composition of the gut microbiota^35-38^.

An alternative treatment for bacterial infectious disease is the use of phages (single or in cocktails), which were first investigated over 100 years ago in the treatment of dysentery^39, 40^. In recent years, phage therapy has gained in popularity, having been effectively used (alone or as adjunctive therapy) to remedy many human infections including chronic rhinosinusitis^41^, urinary tract infection^42-44^, lung and respiratory infection^45-48^, endocarditis^49^, cranial osteomyelitis^50^, and traumatic injury^51^ involving one or multiple bacterial species.

The integration of phages into *in vitro* gut models has provided greater insight into host-phage interactions and the dynamics of the gut microbiota. An *E. coli* phage^52^ and a four-phage cocktail against *C. difficile*^*53*^ were tested in batch fermentation experiments using faecal slurries spiked with host bacteria. Batch models comprised of multiple compartments have been seeded with a seven-strain “ileal microbiota” mix, and tested against *E. coli*^*54*^ and *Listeria monocytogenes*^*55*^ phages. Best representing gut conditions, continuous culture multi-stage systems containing faecal slurries have been utilised to explore *Klebsiella*^*56*^ and *Salmonella*^*57*^ lysis by phages. While some studies also rigorously assessed prophylactic and remedial use^53, 57^, the reports suggested that phages successfully reduced pathogens without impacting the gut microbiota within the system ^52-57^.

One commercially available phage preparation for treating gastrointestinal disease is the Intesti bacteriophage cocktail manufactured by the Eliava Institute (Tbilisi, Georgia), where phage therapy has been in use since the 1920’s^58-60^. This formulation has successfully been used prophylactically and at the onset of disease, and contains in excess of 20 lytic phages for seven pathogenic genera: *Enterococcus, Escherichia, Proteus, Pseudomonas, Salmonella, Shigella* and *Staphylococcus*^58, 61^. Despite its extensive use, safety and efficacy, little is known about how the phage cocktail affects the healthy gut microbiome. In this study, we assessed changes in the adult gut microbiota following Intesti bacteriophage cocktail administration using a human colon model system and shotgun metagenomic sequencing.

## Materials and Methods

### Phage cocktail

The Intesti bacteriophage cocktail was manufactured by the George Eliava Institute of Bacteriophages, Microbiology and Virology in Georgia. The cocktail was purchased over-the-counter at the Eliava Institute, Tbilisi, Georgia for use in these experiments (batch M2-901).

### Media and reaction vessel conditions

Our experiments were conducted using an *in vitro* batch-culture fermentation colon model as described previously ^24^. We used 100 mL reusable glass vessels with a working volume of 50 mL, which were autoclaved prior to experiments. Nutritive media used in the reaction vessel was composed of 2 g/L of each peptone water, sodium bicarbonate (NaHCO_3_) and yeast extract, 0.5 g/L of bile salts and cysteine-HCl, 0.1 g/L of sodium chloride, 0.04 g/L of each di-potassium hydrogen phosphate (K_2_HP0_4_) and potassium di-hydrogen phosphate (KH_2_P0_4_), 0.02 g/L haemin, 0.01 g/L of each magnesium sulphate heptahydrate (MgS0_4_.7H_2_0) and calcium chloride dihydrate (CaCl_2_.2H_2_0), 2 mL of tween-80 and 10 µL of vitamin K1. During the experiments, the temperature was maintained at 37°C with a circulating water jacket and anaerobic conditions were held constant with nitrogen gas bubbling through the media. The pH was held at 6.6 to 7.0 with the addition of 1 M NaOH and HCl solutions using Fermac 260 control units (Electrolab, United Kingdom). A magnetic stirrer at approximately 60 rpm ensured proper mixing of the faecal slurry in the vessels, and the maintenance of anaerobic conditions and pH throughout the experiment.

### Faecal samples and sampling

Three donor faecal samples were obtained from participants recruited onto the Quadram Institute Bioscience Colon Model study. Stool sample collection for use in, *in vitro* fermentation experiments were approved by the Quadram Institute Bioscience Human Research Governance Committee (IFR01/2015) and the Westminster Research Ethics Committee (15/LO/2169), and is registered (http://www.clinicaltrials.gov registration NCT02653001). Informed consent of all subjects was obtained. Participants were over the age of 18, working or living within 10 miles of the Norwich Research Park (coordinates 52.62099801857679, 1.2183262097932304), had normal bowel habits (regular defecation between three times a day to three times a week, with a Bristol Stool type of 3-5), and were not diagnosed with chronic gastrointestinal health problems (such as irritable bowel syndrome, coeliac or inflammatory bowel diseases). At the time of collection, participants completed a survey which confirmed they had not taken antibiotics or probiotics in the previous four weeks, had no gastrointestinal complaints or surgery requiring anaesthetics within the last 72 hours nor were currently pregnant or breastfeeding.

For our study here, three healthy adult donor faecal samples were collected fresh on the morning of the experiment and transported to the institute in triple protected containers at room temperature for immediate processing. Each donor faecal sample seeded two reaction vessels, one for the Intesti bacteriophage cocktail treatment and the other serving as a control. Prior to seeding the reaction vessels, the donor faecal samples were separately homogenised as below. 10 g of the faecal sample was combined in a stomacher bag with 90 mL of sterile phosphate buffered saline at pH 7.4 (PBS, 2.7 mM KCl, 10 mM Na_2_HPO_4_ and 1 mM KH_2_PO_4_). The sample was homogenised with a stomacher 400 (Seward, United Kingdom) for 45 seconds at 230 rpm. 5 mL of the resulting slurry was used to seed 44 mL of nutritive media for both the phage-treated and control vessel. 1 mL of Intesti bacteriophage cocktail was added to the phage-treated vessel 15 minutes after seeding, and 1 mL of PBS to the control vessels. 500 µL samples of the faecal slurries were taken in triplicate from both vessels immediately after the addition of the phage, then again 6 and 24-hours later. The samples were stored at −80°C until further processing.

### DNA extraction and metagenomic sequencing

DNA extraction was performed using two 500 µl aliquots (1 mL total) of the faecal slurries taken at each time point. DNA from 10 mL of the Intesti bacteriophage cocktail and a mock community (suspension of *E. coli* W1485 bacterial cell culture and two lab isolated phages classified as *Jedunavirus* and *Chivirus*) were also extracted to serve as controls for the experiment. Prior to the extraction, bacteriophages and cellular material were co-precipitated by the addition of polyethylene glycol (PEG_8000_) and sodium chloride at final concentrations of 8% and 0.3 M, respectively. The precipitation was incubated at 4°C overnight before being centrifuged at 13,000 ***x g*** for 10 minutes. The supernatant was removed, and the pellet was re-suspended in 750 µL of PowerBead solution (kit supplied) and 60 µL of solution C1 (Qiagen DNeasy PowerLyzer PowerSoil Kit) added. The contents were then subjected to bead beating homogenisation at 4000 rpm for 15 seconds, using a FastPrep-24 classic (MP Biomedicals, USA). The DNA extraction was conducted according to the manufacturer’s instructions (Qiagen DNeasy PowerLyzer PowerSoil Kit). The DNA was eluted in 100 µL solution C6 (kit supplied) and was stored at -80°C prior to sequencing. The DNA samples were sent to Novogene (Cambridge, United Kingdom) for Nextera XT DNA library preparation and shotgun metagenomic sequencing, using the Illumina NovaSeq 6000 system with an S4 flow cell and 150 bp paired end reads.

### Bioinformatics and data analysis

Reads obtained from sequencing were checked for Illumina adapters and the human contaminant and phiX174 sequencing standard reads were removed, and the minimum PHRED quality was set to 20 using fastp^62^. The reads obtained from this study were deposited into the European Nucleotide Archive (ENA) under the study PRJEB96854. For individual sample ENA accessions and associated metadata see supplementary data Table 1. The quality reads were taxonomically profiled with MetaPhlAn3 (v3.0)^63^ and Kraken2 (v2.1.0)^64^ using the k2_pluspf_20210517 database. For more accurate species abundance values, the Kraken2 taxonomic data were re-estimated using Bracken (v2.6.0)^65^. Genera and species relative abundance data were initially visualised in Phantasus (v1.15.4)^66^, and further analysis of the bracken taxonomic data was handled within R (v4.1.2). To compare the profiles obtained and patterns between samples, a heatmap of the relative species (top 500) was generated, using pheatmap (v1.0.12) and viridis (v0.6.4) colour palette packages where values were scaled by row, and the rows were clustered by Euclidean distance and Ward linkage. Further, a barplot of the top six genera (by maximum percentage) was visualised to show abundance changes across all samples. Alpha and beta diversities, analysis of the variance and redundancy were calculated on the species abundance table using the vegan (v2.5-7)^67^ and microbiome (v1.20.0) packages, where the counts were rarefied to the sample with the lowest number of bacterial reads (1.6 x 10^6^; 2B3). The ordination for beta diversity was based on non-metric multi-dimensional scaling (NMDS) of a Bray Curtis dissimilarity matrix, generated by the metaMDS function within the vegan package. Analysis of the community data variance between control and phage treated samples was computed, using the varpart function within the vegan package (on the Bray Curtis dissimilarity) where phage treatment, donor, and time were explanatory variables. A distance matrix and principal coordinates was the basis for a distance-based redundancy analysis (dbRDA) ordination which was constrained by phage treatment.

**Table 1.**
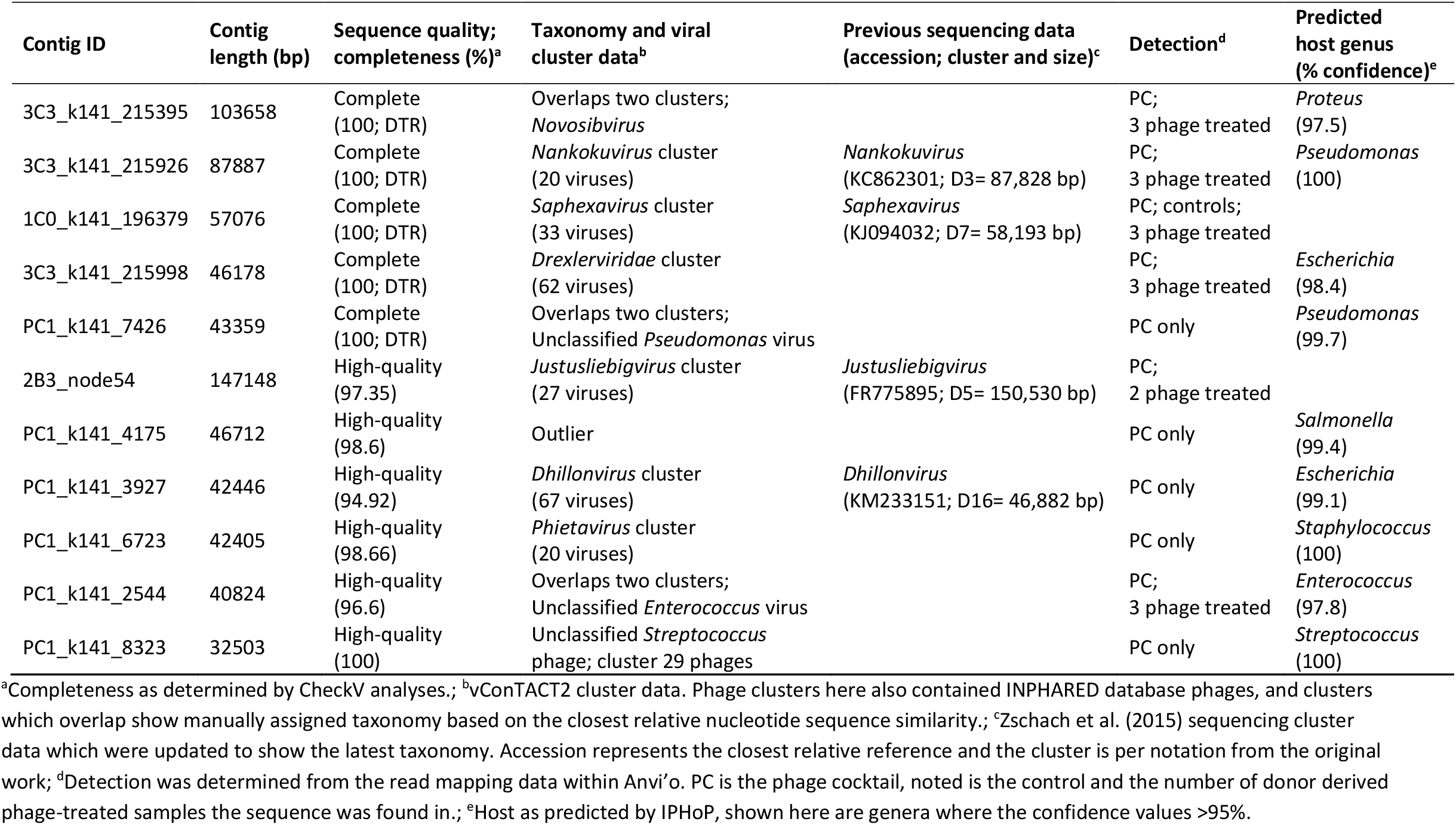
Intesti bacteriophage cocktail sequences.

| Contig ID | Contig length (bp) | Sequence quality; completeness (%)<sup>a</sup> | Taxonomy and viral cluster data<sup>b</sup> | Previous sequencing data (accession; cluster and size)<sup>c</sup> | Detection<sup>d</sup> | Predicted host genus (% confidence)<sup>e</sup> |
| --- | --- | --- | --- | --- | --- | --- |
| 3C3_k141_215395 | 103658 | Complete (100; DTR) | Overlaps two clusters; *Novosibvirus* |  | PC; 3 phage treated | *Proteus* (97.5) |
| 3C3_k141_215926 | 87887 | Complete (100; DTR) | *Nankokuvirus* cluster (20 viruses) | *Nankokuvirus* (KC862301; D3= 87,828 bp) | PC; 3 phage treated | *Pseudomonas* (100) |
| 1C0_k141_196379 | 57076 | Complete (100; DTR) | *Saphexavirus* cluster (33 viruses) | *Saphexavirus* (KJ094032; D7= 58,193 bp) | PC; controls; 3 phage treated |  |
| 3C3_k141_215998 | 46178 | Complete (100; DTR) | *Drexleriviridae* cluster (62 viruses) |  | PC; 3 phage treated | *Escherichia* (98.4) |
| PC1_k141_7426 | 43359 | Complete (100; DTR) | Overlaps two clusters; Unclassified *Pseudomonas* virus |  | PC only | *Pseudomonas* (99.7) |
| 2B3_node54 | 147148 | High-quality (97.35) | *Justusliebigvirus* cluster (27 viruses) | *Justusliebigvirus* (FR775895; D5= 150,530 bp) | PC; 2 phage treated |  |
| PC1_k141_4175 | 46712 | High-quality (98.6) | Outlier |  | PC only | *Salmonella* (99.4) |
| PC1_k141_3927 | 42446 | High-quality (94.92) | *Dhillonvirus* cluster (67 viruses) | *Dhillonvirus* (KM233151; D16= 46,882 bp) | PC only | *Escherichia* (99.1) |
| PC1_k141_6723 | 42405 | High-quality (98.66) | *Phietavirus* cluster (20 viruses) |  | PC only | *Staphylococcus* (100) |
| PC1_k141_2544 | 40824 | High-quality (96.6) | Overlaps two clusters; Unclassified *Enterococcus* virus |  | PC; 3 phage treated | *Enterococcus* (97.8) |
| PC1_k141_8323 | 32503 | High-quality (100) | Unclassified *Streptococcus* phage; cluster 29 phages |  | PC only | *Streptococcus* (100) |

<sup>a</sup>Completeness as determined by CheckV analyses.; <sup>b</sup>vConTACT2 cluster data. Phage clusters here also contained INPHARED database phages, and clusters which overlap show manually assigned taxonomy based on the closest relative nucleotide sequence similarity.; <sup>c</sup>Zschach et al. (2015) sequencing cluster data which were updated to show the latest taxonomy. Accession represents the closest relative reference and the cluster is per notation from the original work; <sup>d</sup>Detection was determined from the read mapping data within Anvi'o. PC is the phage cocktail, noted is the control and the number of donor derived phage-treated samples the sequence was found in.; <sup>e</sup>Host as predicted by IPHoP, shown here are genera where the confidence values >95%.

To best recover viruses from the samples, we utilised an assembly approach where the high-quality reads were assembled with MEGAHIT (v1.1.3)^68^ using default k-mers lengths of 21,29,39,59,79,99,119,141. Viral and phage sequences were predicted from these assembled contigs with VIBRANT (v1.2.1)^10^ using the default settings with a minimum sequence length of 1 Kb and VirSorter2 (v2.2.3)^12^ with a minimum sequence length of 1.5 Kb and the default dsDNA/ssDNA database. VIBRANT and VirSorter2 predicted viral sequence files (for each individual sample) were labelled and concatenated with SeqFu (v1.9.1)^69^. This resulted in a collection of uncultivated viral genomes (UViG) predicted for all the samples which could be prepared for further analyses using Anvi’o (v7.1)^70, 71^. The individual UViG files (.fasta) were concatenated with SeqFu, twice dereplicated firstly using CD-HIT-EST (v4.8.1)^72^ at 95% identity (-c 0.95) with a 99% sequence alignment (-aS 0.99) and then by CheckV (v1.0.1)^73^ rapid genome clustering with average nucleotide identity of 95% and an alignment fraction of 85%, to approximate species-level clusters for dsDNA prokaryotic viruses of the class *Caudoviricetes* as defined by the International Committee on Taxonomy of Viruses (ICTV)^74^. The dereplicated dataset consisted of species level vOTUs (viral Operational Taxonomic Units) which were each represented by their largest UViG. Only the vOTUs greater than 4.5 Kb were retained, resulting in the final contig/vOTU database. Coverage and detection of the vOTUs from the contig database were established through read mapping of the donor and phage cocktail read libraries with Bowtie2 (v2.4.1)^75^ end-to-end mapping (default settings and penalties) in conjunction with SAMtools (v1.12)^76^. Within Anvi’o (v7.1), the resulting BAM files were individually profiled against the contig database which was functionally annotated using the NCBI COGs database (COG20)^77^, and had tRNAs identified^78^. The profiles were then merged to observe patterns in the viral species present. Any vOTUs showing mapping with the mock community were excluded from further analysis.

The taxonomic diversity of the phage sequences found in the contig database were assessed using a gene-sharing network with vConTACT2^79^, annotated using the INPHARED pipeline^80^. Briefly, the contigs in the vOTU database were annotated using PHANOTATE with the Pharokka pipeline (v1.1.0)^81^. The annotated files were transformed into the input format for vConTACT2 using the script single_GenBank_to_vConTACT_inputs (https://github.com/RyanCook94/Random-Perl-Scripts/blob/main/single_GenBank_to_vConTACT_inputs.pl). The annotated contig .faa file and the INPHARED database file 2Nov2022_vConTACT2_proteins.faa were concatenated and used as input for the --raw-proteins in vConTACT2, as were the gene to genome mapping files for input as the -- proteins-fp. vConTACT2 was run without an internal database, using diamond^82^ for protein clustering and ClusterONE^83^ for genome clustering. The resulting network was visualised with Cytoscape (v3.9.1)^84^ and annotated with the INPHARED database annotation tables dated 2 November 2022.

Where comparative genomics between our phage and database phages was required, we annotated individual genomes (as above using PHANOTATE in the Pharokka pipeline v1.1.0) before gene clusters were analysed using Clinker (v.0.0.23)^85^. Individual read mapping across one genome was viewed by Integrative Genomics Viewer (IGV; v2.6.1)^86^. Input files for IGV were generated with SAM tools (v1.12) and BamToCov (v2.7.0)^87^.

To link potential phage sequences with their predicted bacterial hosts we used iPHoP (v1.2.0)^88^. Phage lifestyle was predicted as a part of CheckV “prophage” analysis and by BACPHLIP (v0.9.3)^89^. Potential prophage activity was estimated with PropagAtE (v1.0.0)^90^ using MEGAHIT assembled contigs and VIBRANT generated prophage coordinates.

## Results

To simulate the human gut system and gut microbiota, we used an *in vitro* batch-culture fermentation colon model seeded with faecal samples from three healthy donors. For assessment of the 5microbial changes, each donor had an untreated control and a phage cocktail treated vessel, and these faecal slurries were sampled over time at phage addition (0 hours), 6-hours and 24-hours. The total microbial DNA was extracted and sequenced, and we obtained 21-35 million quality reads per sample (Supplementary Table 1).

### Composition of the microbiome

Across the three donor samples, we detected over 100 distinct bacterial genera and viral species with scantly detected archaeal and no fungal signatures. At the species level, little overlap in gut microbial composition was observed between individuals (Figure 1A). While the exact composition was unique, several bacterial genera which are typical members of the human gut microbiota including *Bacillus, Bacteroides, Eubacterium, Faecalibacterium, Lactobacillus* and *Phocaeicola* were detected in all three donors. *Bacteriodes* and *Faecalibacterium* were ubiquitous, detected in the top six most abundant genera of both the control and phage treated samples (Figure 1B). Whereas genera such as *Bifidobacterium, Prevotella* and *Streptococcus* were more abundant in specific donors (donor 1, 3 and 2 respectively) and their abundance decreased over time.

**Figure 1.**
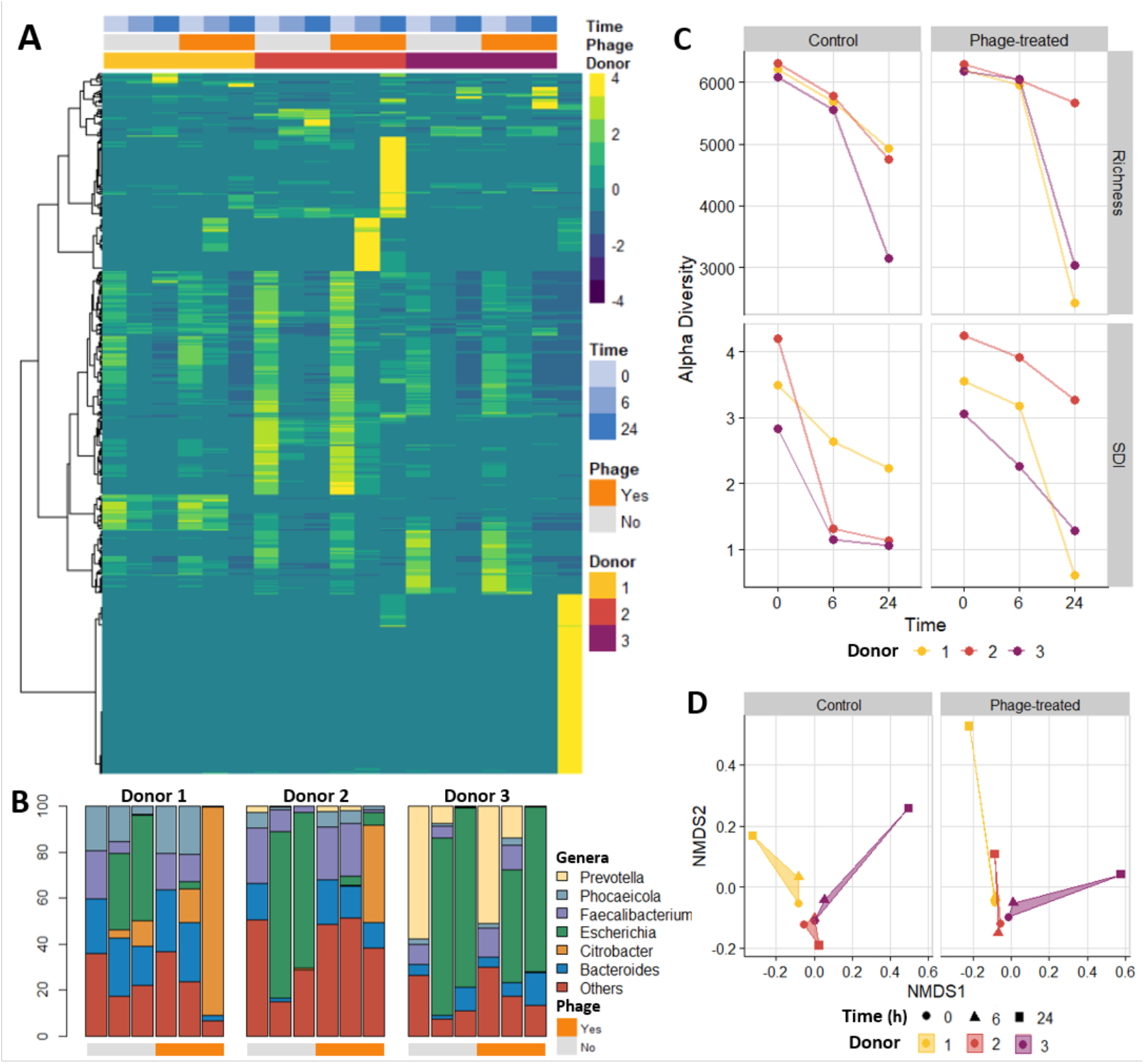
Microbial diversity of samples. **(A)** Heatmap of the relative abundance of the top 500 species as determined by Bracken. The columns represent the samples and the phage cocktail (rightmost column). The rows indicate the identified viral or bacterial species, and these were hierarchically clustered by Euclidean distance and Ward linkage. The relative abundance was scaled by row. **(B)** Genus relative abundances across samples. Samples are grouped by donor, ordered by time (0, 6 and 24 hours), and are divided by treatment where control and phage treatment are coloured by grey and orange respectively. **(C)** Alpha diversity indices of the samples (Richness and Shannon Diversity Index [SDI]) separated by phage treatment. Points represent individual samples across time points. **(D)** Sample beta diversity and NMDS plots according to phage treatment. The stress value for the plot is 0.26.

Initially, donor one derived samples contained similar proportions of *Bacteriodes, Faecalibacterium* and *Phocaeicola* (Figure 1B), with the remaining portion composed of *Ruminococcus, Bifidobacterium*, and many other genera. Whereas the samples derived from donor two had more *Faecalibacterium* present with less *Bacteroides*, and all other genera were below 10% relative abundance each. *Prevotella* dominated samples derived from donor three, with a relative abundance over 50% (Figure 1B). These initial compositions altered markedly over the course of the experiment (Figure 1B). Opportunistic and adaptive bacteria belonging to the genera *Escherichia, Klebsiella, Proteus, Salmonella, Serratia* and *Veillonella* had the highest abundance after the initial time point in one or more donor derived samples. This was particularly evident for *Escherichia* and *Citrobacter* (Figure 1B). As the microbiota adapted to the condition of the model gut system, we observed reductions in the alpha diversity (Figure 1C). The richness and Shannon Diversity Index (SDI) were the highest initially then decreased over the course of the experiment, a trend seen in both the control and phage treated samples (Figure 1C). Likewise, the beta diversity at first was similar across samples before diverging over time (Figure 1D).

These analyses revealed changes in bacterial composition not only with phage treatment, but by individual donor, and time. To estimate how much of the community variation could be attributed to each variable we used Bray Curtis dissimilarity in conjunction with variation partitioning. Changes over time contributed 31.5% of the variance in the composition data, and donors represented 14.2%, whereas phage treatment only contributed 2.8% of the variance (adjusted R^2^ values). Next, we used distance-based redundancy analysis (dbRDA) to identify the main changes in bacterial relative abundance according to phage treatment (Figure 2A). The bacterial species most impacted was *Escherichia coli*, which was more closely associated with the controls. In contrast, we observed that *Citrobacter koseri* and *Faecalibacterium prausnitzii* were more abundant in phage treated samples, whereas *Prevotella copri* showed similar relative abundances, irrespective of the phage treatment (Figure 2A). The relative abundance of these species differed by time and donor (Figure 2B-E). *E. coli* abundance increased after the initial timepoint, was detected in all donor derived samples, and was reduced in phage treated samples compared to the controls (Figure 2B). Similarly, *C. koseri* abundance increased over time, however, the species was more prevalent in the phage treated samples of two donors, especially those derived from donor 1 (Figure 2D). Over the course of the experiment, *P. copri* abundance decreased in donor 3 control and phage treated samples (Figure 2C). *F. prausnitzii* relative abundance also decreased over time, but was found in all three donor samples (Figure 2E). While full interactions and population dynamics in the system were not known and our sample size did not permit statistical testing, our observations suggest that *E. coli* abundance was reduced by the lytic action of phages from the cocktail. Lysis due to the phage cocktail phages also could have left nutrients for other bacteria such as *F. prausnitzii* to grow within the model.

**Figure 2.**
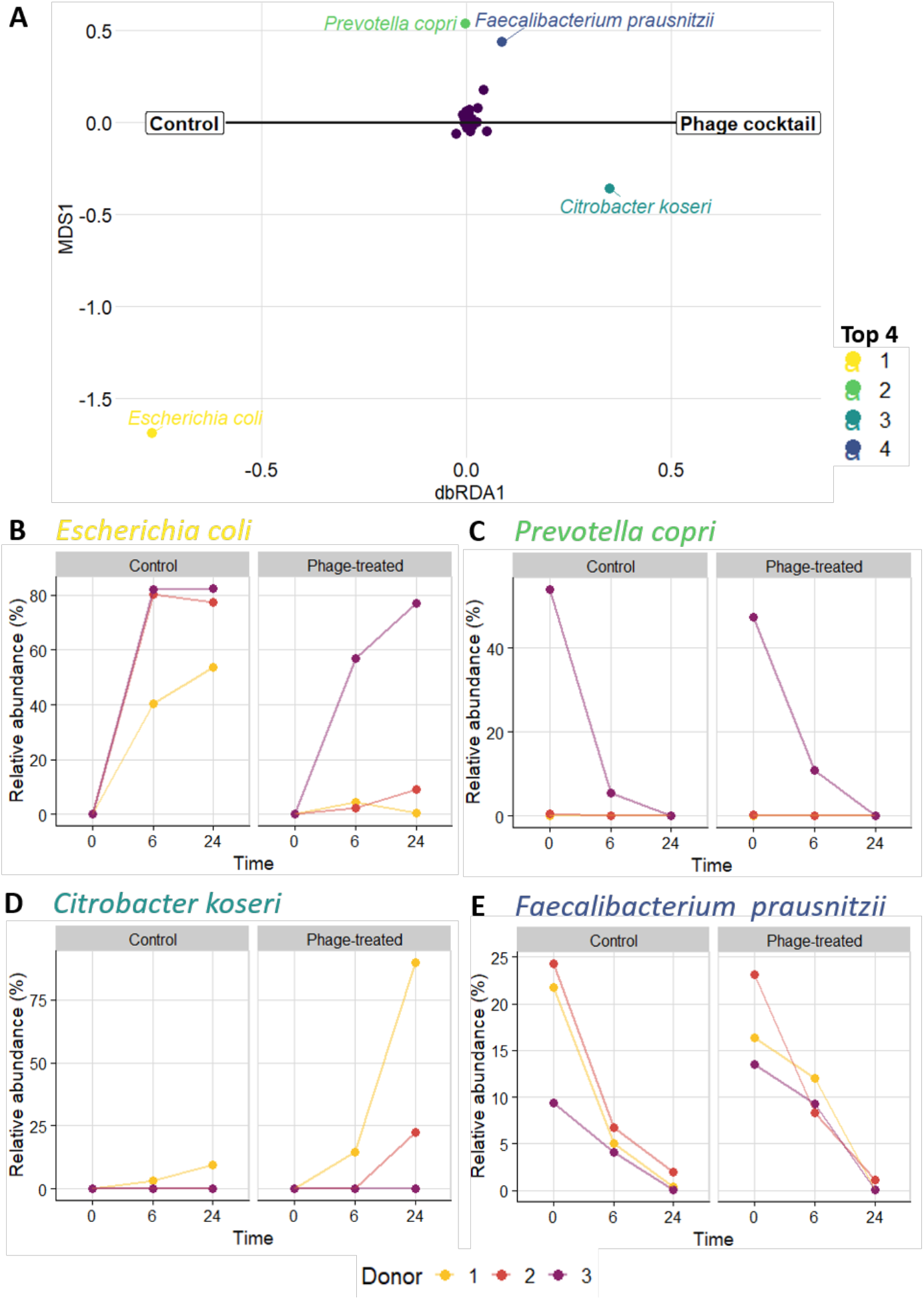
Bacterial community composition dynamics separated by phage treatment. A distance-based redundancy analysis (dbRDA) plot (**A**) constrained by phage treatment (X axis), showing the top four most impacted bacterial species. Their location relative to the centerpoint (X=0) indicates association with control (left) or phage treated (right) samples. Plots of the relative abundance of the top four species (**B-E**), separated by phage treatment, illustrate variation over time and by donor.

### Phage cocktail phages have the potential to actively replicate in the healthy gut

We next looked at the phage portion of the microbiome, to investigate whether phages from the cocktail were actively replicating in our model system, thereby influencing the bacterial composition. This analysis was complicated by the observation that the read profilers were not able to distinguish well between different phage signatures, especially at the species level. We, therefore, used the gold-standard pipeline of assembly, phage identification, viral Operational Taxonomic Unit (vOTU) clustering and read mapping for analysis (see methods; as previously described^91^). Across all libraries excluding the mock community, we generated a virus contig database of 5306 vOTUs where each vOTU was represented by the largest contig or Uncultivated Virus Genome (UViG), which were classified by CheckV as complete (66/5306), high-quality (103/5306), medium-quality (182/5306), Low-quality (2189/5306) and the remainder were not-determined (2766) (Supplementary Table 2; vOTU database available on FigShare). Read mapping of these vOTUs and visualisation with Anvi’o, allowed us to discern which vOTUs were present in the phage cocktail. These analyses revealed that only 166 vOTUs recruited reads from the phage cocktail library using a detection limit of 0.25 (i.e. minimum 25% of the contig length covered by reads) and minimum average coverage over one (Figure 3). It is important to note here that we chose this approach because the phage cocktail itself had a low number of phages per mL and direct assembly yielded only a low number of high-quality UViGs.

**Figure 3.**
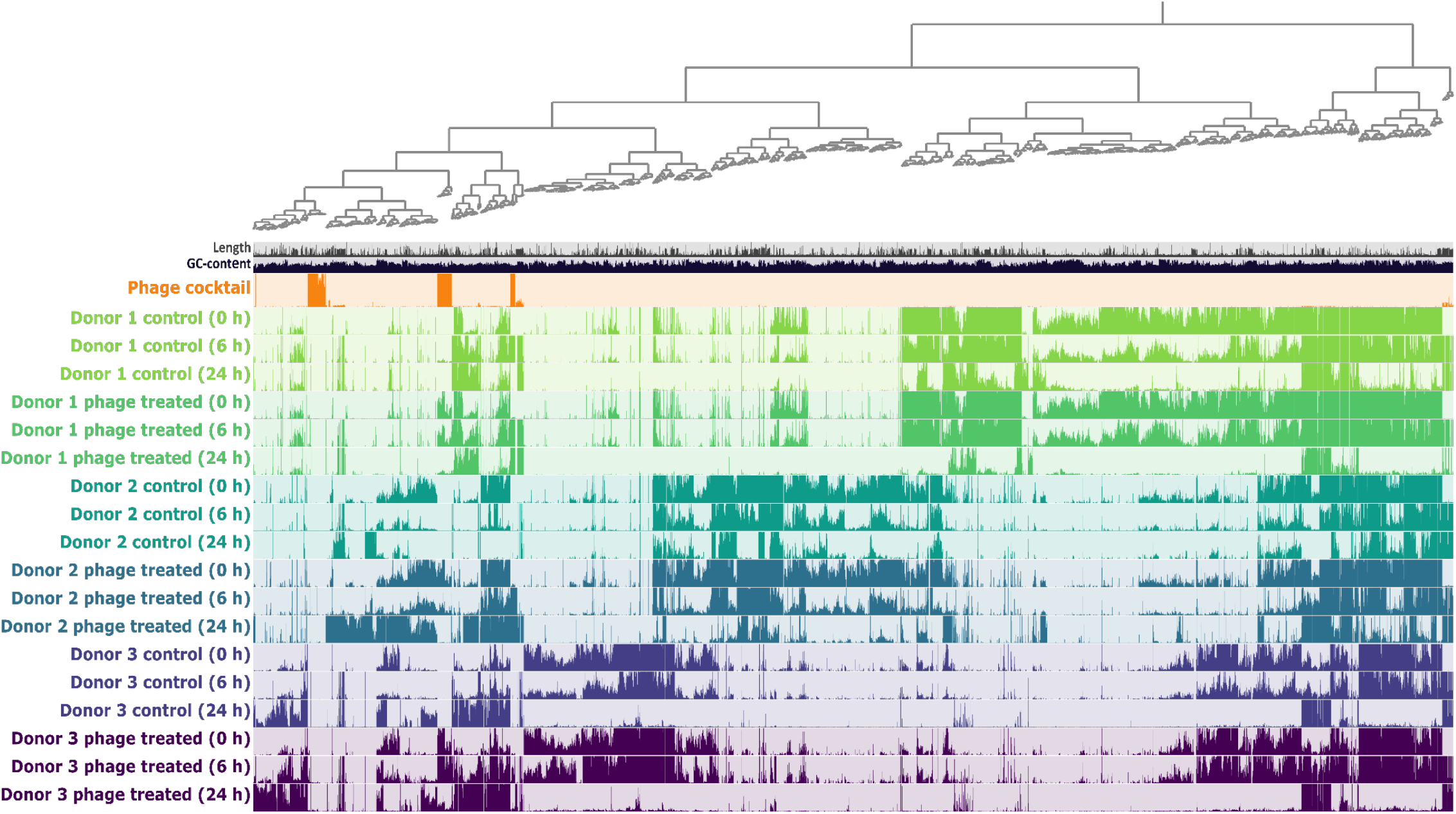
Differential abundance of the virus contig database. Tracks represent sample sequence libraries and each column a vOTU or split (contigs were split in multiple sections if length was longer than 20 kb). The height of the bar within the tracks depicts mean coverage of vOTUs or splits to a maximum of 10x coverage. The dendrogram shows clustering according to sequence composition and differential coverage (Euclidean distance and Ward linkage). Figure created using Anvi’o

The percentage of mapped reads from each sample recruited to the phage cocktail vOTUs, ranged from 76% (donor 2 samples at six hours) to 0.1% (as seen in all controls at time zero), with an average depth of coverage from 1292X to almost zero (Supplementary Table 3). The phage cocktail vOTUs ranged in size from 4.5 to 147 kb, with approximately half over 10 kb in length. Direct terminal repeats (DTRs) were identified in five sequences which were marked as complete by CheckV and of the remaining, six were classified as high-quality, six as medium-quality and 149 as low-quality (Supplementary Table 2). We discovered that 75 of the 166 vOTUs detected were exclusively found in the cocktail, and conversely 24 were detected in phage treated samples as well as control samples, indicating that a subset of phages in the phage cocktail are related to natural members of the gut microbiota. Of the 67 vOTUs found only in phage treated vessels, the majority (46/67) were detected in all phage treated vessels suggesting detection irrespective of their replication within the model, whereas others found in one (18/67) or two (3/67) donor derived samples may be reliant on replication for detection.

To gain insight into the taxonomic diversity of the vOTUs found in the phage cocktail, we constructed a gene sharing network using vConTACT2, where each phage or vOTU contig is a node connected with an edge if they share a significant number of genes. The network revealed that vOTUs detected in the phage cocktail were more closely associated to reference sequences as opposed to donor derived viral sequences detected in our study (Figure 4). Further investigation of the 166 phage cocktail vOTUs showed that 60 were assigned viral clusters and of these only 14 also contained representatives from the INPHARED database. An additional five vOTUs were found to overlap clusters, while the number of singletons and outliers totalled 101 (Supplementary Table 2 and 4). Considering the clustering, read mapping patterns and completeness, our sequencing here produced 11 unique phage genomes which were marked (by CheckV analysis) as complete with DTRs or high-quality genomes from the phage cocktail (Table 1). We also observed the presence of partial phage genomes which pertained to six unique viral clusters containing database representatives, and numerous other contigs which were singletons or outliers in the gene sharing network, and of low quality (Supplementary Table 2 and 4). The phage cocktail contained sequences from diverse virus taxa, clustering with known members from *Dhillonvirus, Justusliebigvirus, Nankokuvirus, Phietavirus* and *Saphexavirus* genera (Figure 4; Table 1). Of note, we also identified a high quality UViG which clustered with 29 unclassified *Streptococcus* phages, a genus or host not reported to be targeted by the Intesti bacterophage cocktail. Further diversity was discovered in shorter vOTUs clustered with known larger viral sequences from genera such as *Astrithrvirus, Berlinvirus, Chivirus, Peduovirus* and *Tequintavirus* (Figure 4; Supplementary Table 2 and 4), as well as outlier and singleton BLASTn similarity (at species level demarcation >95% identity over 99% query) with *Asteriusvirus, Kayvirus, Felixounavirus, Kochikohdavirus, Mosigvirus and Tequatrovirus*.

**Figure 4.**
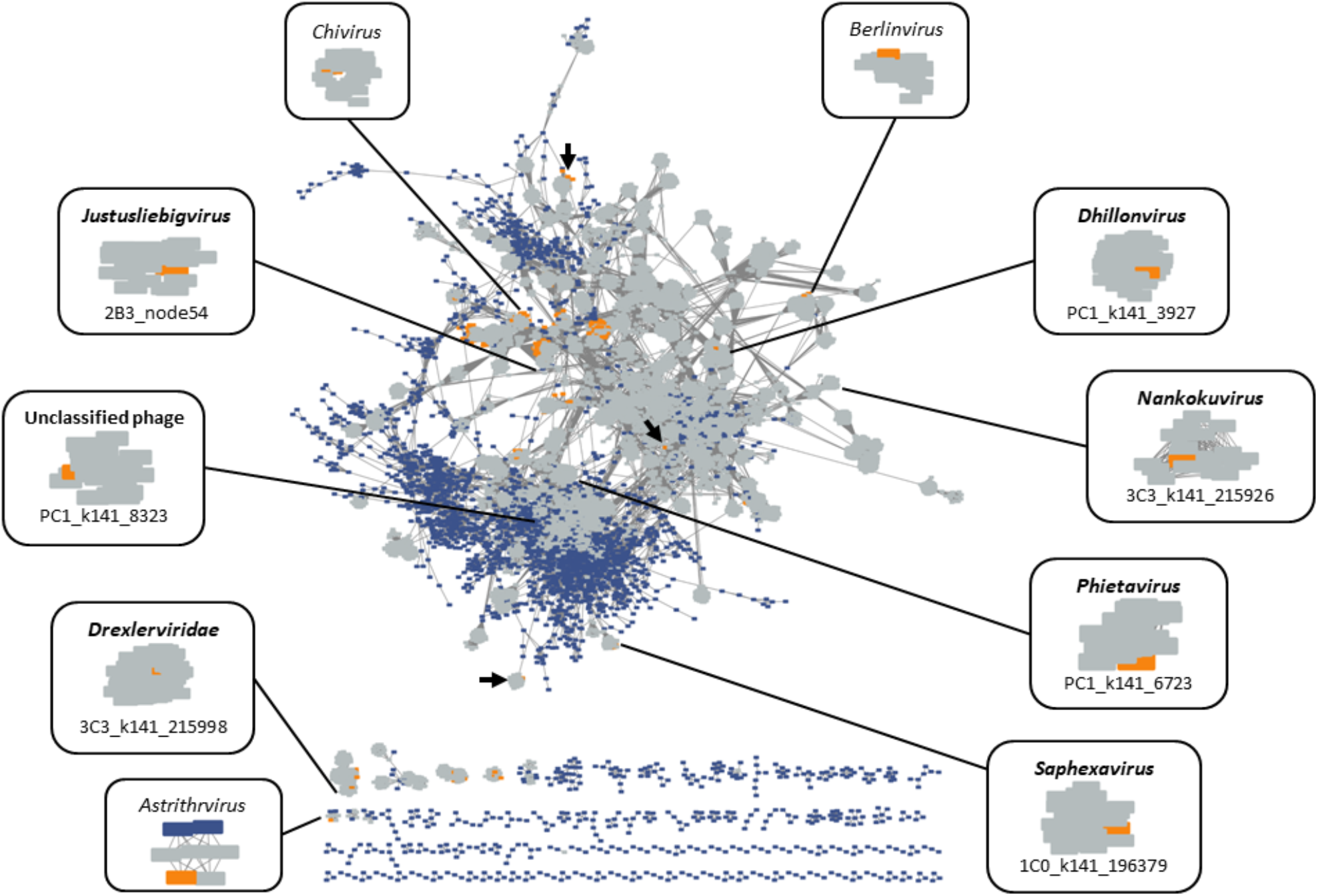
Predicted vOTU Gene-sharing network. Known phages from the INPHARED reference database are represented as coloured grey rectangles, all contig database predicted vOTUs across our dataset are marked in blue and vOTUs detected in the phage cocktail are marked in orange. Annotated are virus genera or families, and individual phage which cluster at that classification. Complete or high-quality sequences which overlap clusters are marked with arrows

We compared our clustered UViGs to previously defined phage clusters by Zschach et al. (2015) to explore the diversity of phages within the Intesti bacteriophage cocktail, and discovered that only four (out of 11; 36%) of our UViGs group with the 18 previously described phage clusters (Table 1) and an additional two were clustered with incomplete vOTUs from our data (*Berlinvirus* and *Vequintavirus*). While not clustered, some vOTUs representative of the *Chivirus, Drexlerviridae* and *Kayvirus* genera were identified in both studies. Our sequencing here revealed increased diversity from the phage cocktail, as we identified UViGs which cluster with known *Phietavirus, Novosibvirus* and an unclassified *Enterococcus* phage, as well as partial vOTUs from *Peduovirus* and *Kochikohdavirus*. Furthermore, host prediction indicated that these phages could target members of *Enterococcus, Escherichia, Proteus, Pseudomonas, Salmonella, Staphylococcus* and *Streptococcus*, and therefore, could impact the microbiota in our gut model (Table 1).

As phages may influence the bacteria, we looked for phages undergoing active infection in our model which we regarded as phage cocktail vOTUs with increased read mapping at 6 and 24-hours post inoculation, and reads mapped to at least 80% of the vOTU length (detection limit of 0.8). We identified a subset of 32 vOTUs which after six hours had average coverage over 10X, however, these often were not representative of full genomes, not clustered with known INPHARED database phages, nor had a predicted host (Supplementary Table 3 and 5). Their elevated coverage indicated growth in the model system but their impact on the community was unknown.

Considering the UViGs identified in our study, six were also detected in selected donor derived samples and were elevated compared to the control (Table 1; Supplementary Table 3). In donor 3 derived samples, we discovered that the *Novosibvirus, Nankokuvirus, Saphexavirus, Drexlerviridae* (family) and unclassified Enterococcus phages had the highest read mapping (<100 times) at six hours, demonstrating the phages response to fluctuations in their host bacteria over the course of the experiment. We uncovered the largest increase in average coverage across all phage cocktail vOTUs from the high-quality 147 kb *Justusliebigvirus* phage (Table 1) which had closest sequence similarity and gene structure to Enterobacteria phage phi92 with 247 CDS and 14 tRNA (Figure 5A). This phage, we labelled as “justusliebigvirus bloom phage”, was overrepresented in donor 2 derived samples at six and 24-hours with average coverage of over 23,000X and 1700X respectively, as well as approximately 80X coverage in the donor 3 derived sample at 24 hours (Figure 5B). We found this phage at a low average coverage (1.45X) in cocktail and other donor derived samples (Figure 5B), thus replication must have occurred within the vessel and this likely depended on the growth of *Escherichia coli*.

**Figure 5.**
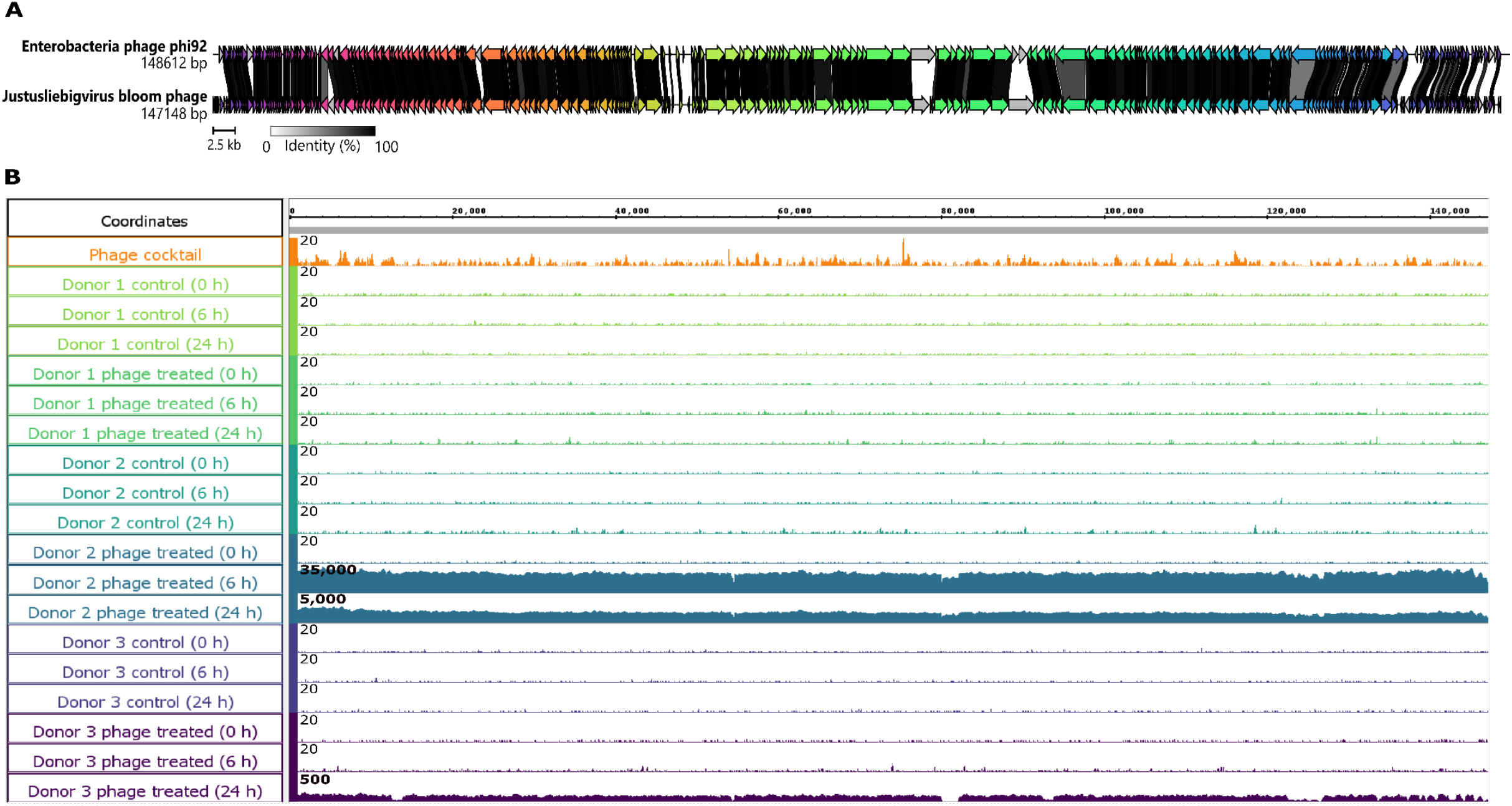
Genome representation of the phage present in the phage bloom and read mapping across the donor derived samples. Represented is the genome structure and comparative genomics between the Justusliebigvirus bloom phage and Enterobacteria phage phi92 (A) where arrows indicate predicted genes, colour shows homologous genes and the links nucleotide similarity between the coding sequences. Read mapping of the genome with donor derived sample libraries (B). Each track represents a sample and the height of the bars represent the reads mapped at that location with the maximum of the scale noted at the upper left corner.

Also present with high coverage at the same time point and vessel as the justusliebigvirus bloom phage were several shorter vOTUs. While these vOTUs did not cluster in the vConTACT2 network, they shared similarity to multiple members of the genus *Tequatrovirus* by BLASTn similarity at species level demarcation and we hypothesise that these contigs together could be equivalent to a predicted complete genome which did not assemble. These two phages together (justusliebigvirus bloom phage and unknown *Tequatrovirus*) account for approximately 85% of the viral reads in the donor 2 derived six-hour sample.

### Impacts of the phage cocktail on the gut microbiota are often transient

Given the replication of the justusliebigvirus bloom phage and the apparent lysis of *E. coli* within the donor 2 phage treated vessel, we searched for subsequent effects by scanning for elevated vOTUs at 24-hours. From the entire phage contig database, we uncovered 12 high quality vOTUs that had their highest average coverage at this time point, 10 of which were donor 2 specific, and three clustered within our vConTACT2 network (Supplementary Table 6 and 7). A total of eight vOTUs were only detected at 24-hours, including one (2B5_k141_64032) that clustered with unclassified Escherichia phage vB_EcoM-613R2, a prophage of *E. coli* ^92^, and was marked by BACPHLIP as temperate (Supplementary Table 6). Also clustered with *Rauchvirus* prophages was another vOTU (2B5_k141_55335) which was detected initially in both the control and phage treated samples, and then again at 24-hours but only in the phage treated vessel with over a hundred times average coverage. These two vOTU likely would have relied on the stress of *E. coli* and *Bordetella* respectively which occurred prior to 24-hour sample. The final vOTU to cluster (with three unclassified phages), 2B5_k141_91452, was marked as lytic but was only present at the final time point in excess of 170 times coverage in the phage treated vessel but minimally in the control. The known phages in the cluster all have unique hosts, and a host was not predicted for the vOTU with iPHoP, but it is likely that the host was able to make use of biproducts produced from other bacteria or was non-fastidious. The impact of this vOTU on the community and those also unclassified was unknown.

Next, we investigated whether typical members of the gut virome were altered in the phage cocktail treated vessels, by observing changes in average coverage of vOTUs clustered with members of the *Crassvirales* order and the *Microviridae* family. From our vConTACT2 network, we identified three *Crassvirales* and two *Microviridae* vOTUs which were marked as lytic by BACPHLIP (Supplementary Table 8). The three *Crassvirales* vOTUs were representative of three families: *Crevaviridae, Intestiviridae* and *Steigviridae*, and were specific to donor 3, 2 and 1 derived samples respectively, within the control and phage treated vessels. The *Crevaviridae* vOTU (3A0_k141_46950) in the phage treated vessel had coverage approximately four-times higher average coverage than the control at six hours, however, at 24-hours the level was equivalent to the control. Although shorter than a predicted genome, we uncovered that the *Intestiviridae* vOTU (2B5_k141_39930) had a 100-fold increase in coverage compared with the control at 24-hours. Conversely, the *Steigviridae* (1A0_k141_190349) vOTU showed similar patterns of coverage across six and 24-hours. We discovered that the two *Microviridae* vOTUs were exclusive to donor 2 control and phage treated samples. One vOTU (2A0_k141_222313) was maintained across timepoints, whereas the other (2B0_k141_198670) showed the highest coverage at the initial timepoint, but both were concordant with the control (Supplementary Table 7 and 8).

Our analysis of the bacterial community composition suggested that there were increased levels of *Citrobacter koseri* and *Faecalibacterium prausnitzii* signatures detected in the presence of the phage cocktail. In effort to link these bacteria with vOTU abundance changes, we searched for those clustering with known phages, and subsequently alterations in their coverage. There were no *Citrobacter* phages clustered with the vOTUs in our database, however, guided by the Kraken2 results, we discovered the top six vOTU (by average coverage) in the donor 1 phage treated vessel at 24-hours, had BLASTn similarity (at species demarcation) to *C. koseri* (Supplementary Table 7 and 9). With thousands of times coverage, at levels ten times greater than the control, two vOTU (1A5_k141_68270_0; 2B5_k141_36022) indicated stress of the bacteria and bacterial defence systems. As one was a 19 kb fragment that contained a portal protein, and may indicate release of a cryptic prophage or one which did not assemble, and the other had a type I-F CRISPR/Cas system (contained within an 86 kb sequence) with two short CRISPR loci that may have prevented phage predation of the bacteria. While the remaining vOTUs were marked as bacterial (false positives) or were detected in the mock community (possible contaminants), one (1A5_k141_30475), a 300 kb vOTU, contained a 50 kb predicted complete prophage sequence (as determined by checkV) at levels 10-fold greater than the control. Further analysis showed that mock community reads mapped to bacterial genes outside of the predicted prophage region, and moreover indicated there were multiple prophages at this time point, that may have contributed to the levels of *Citrobacter* detected within the vessel. We identified in the dataset, six high-quality and complete vOTUs that clustered with *Faecalibacterium* phages, but their coverage changes were small and we could not link these to bacterial abundance (Supplementary Table 7 and 10). A result also reflected in the vOTUs which were predicted to have *F. prausnitzii* as a host.

One potential side effect of phage cocktail treatment could be the induction of prophages from the native bacteria present in the donor sample. To investigate whether the temperate phages identified in our gut model were actively replicating, PropagAtE prediction was used, where differences in read mapping to the bacterial chromosome and the prophage coordinates inferred replication status. With this tool, the number of induced prophages found in all samples were low and there were no apparent differences between the phage treated and control samples (Supplementary Table 11). However, individual phage-bacteria interactions by induced prophages such as those hypothesised in donor 2 derived samples (as detailed above), and other episomal prophages interactions could have been impacting the bacterial community over time.

## Discussion

An ideal phage cocktail is one that contains strictly lytic phages which lyse pathogenic bacteria contributing to disease, while leaving the resident microflora unaffected. Given the complex microbial structure of the human gut and its integral role in health^1^, effective and specific lytic phages are particularly advantageous. Phage preparations such as the Intesti bacteriophage cocktail are available for purchase to treat common gastrointestinal complaints, and when taken orally are exposed to the resident gut microbiota. However, the impact on these communities is understudied. Our study here described changes in both the bacterial and viral communities from three healthy individuals in the presence of the Intesti bacteriophage cocktail, using a simulated gut model. Despite the many complex dynamics and interplays, our results suggest that fluctuations in bacteria and viruses associated with the phage cocktail had minimal interruption to the native gut microbiota within the model.

The gut models linked with all three donors were colonised by the typical members of human gut microbiota. We detected the presence of *Bacteriodes* and *Faecalibacterium* in all donor samples whereas genera such as *Bifidobacterium, Prevotella* and *Streptococcus* were donor specific and often transient across time points, which is consistent with community dynamics in the gut ecosystem^93^. Our profiling and the community composition at the initial time point indicated the donors had healthy gut microbiome enterosignatures, where two microbiomes were *Bacteriodes* (ES-Bact) and the other was *Prevotella* (ES-Prev) centric. Each donor had a unique bacterial community at the species level and the diversity decreased over the course of the experiment. This was reflected in the calculated diversity indices and increased abundance of non-fastidious bacteria such as *Escherichia*. Given the static nature of our experimental model and once-supplied media within the system, this decrease in diversity was expected and was a limitation of the study.

The bacterial community composition altered over the course of the experiment. Our analysis using time, donor and phage treatment as explanatory variables and Bray-Curtis distance revealed the largest impact on the community variance was time, whereas phage treatment contributed under 3%. When comparing phage treatment to controls, the most impacted species was *E. coli* which was less abundant in phage treated samples at 6 and 24 hours. Although our study is not powered for statistical testing, we observed a reduction of *E. coli* signatures in all phage treated vessels, and detected *E. coli* phages within the experimental system (as discussed below) which potentially impacted the bacterial community. While the full pathogenicity and impact of the strains remains unknown, a decreased abundance of *E. coli* may be beneficial in a treatment setting to reduce disease symptoms and restore gut eubiosis.

Our sequencing effort here captured as yet uncultured phage isolates, as compared with the previously reported cocktail diversity, and indicated that the Intesti bacteriophage cocktail is composed of numerous phages from various taxonomic genera. Previous work by Zschach et al. (2015) reported the cocktail to contain 22 phage clusters which were derived by propagating phages from the preparation. Our direct sequencing approach, creation of a contig database followed by read mapping and vOTU clustering, revealed both taxonomically similar and diverse phages within the phage cocktail at the genus level, and potentially novel taxonomic diversity. The presence of an unclassified *Streptococcus* phage with an average coverage of 17X over the entire sequence from the phage cocktail library, was unexpected as it is not a host reported to be targeted by the Intesti phage cocktail. This phage was not found to replicate in phage treated vessels and may have been introduced to the cocktail prior to our study.

Productive infection of Intesti cocktail-derived was occurring within our gut model. We observed the justusliebigvirus bloom phage, which was overrepresented in donor 2 derived samples with average coverage in excess of 1000 times. Its high sequence identity to the *E. coli*-infecting phage phi92, makes it likely that the bloom phage also infects *E. coli* or a related strain. Similarly, most members of the genus *Tequatrovirus* are known to lyse *E. coli* or *Shigella* strains, and we therefore speculate that this minor bloom phage also infects *E. coli*. In parallel, we observed a decrease in *E. coli* signatures, not just in the sample undergoing the active phage bloom, but also in other phage-treated donors and time points. From this, we have good evidence to conclude that the Intesti bacteriophage cocktail influences the *Escherichia* populations in the gut. However, we currently do not have enough sequence data to ascertain what strains of *E. coli* are affected, and whether these are considered commensals in healthy adults or whether they are opportunistic pathogens that could lead to disease development.

The microbiota in the system adapted to changed conditions. Following the phage bloom in the donor two phage treated vessel, there was an increase in high quality vOTUs at 24 hours. The majority of these were specific to the donor 2 vessel at that time point, and include both lytic and temperate vOTU (as predicted by BACPHLIP). A vOTU which clustered with *Rauchvirus* prophage was detected initially and again at 24-hours, indicating the potential for native donor phages to modulate gut bacteria.

Phage treatment appeared to minimally affect the native microbiota. Typical members of the gut virome, *Crassvirales* and *Microvidae* viruses were detected in our dataset but each vOTU was limited to one donor’s sample, rather than being ubiquitous. We observed that vOTUs were often present across all time points and concordant in both control and phage treated samples, indicating their stability and persistence in the donor microbiota. Our analysis showed an increase of *C. koseri* and *F. prausnitzii* in phage treated vessels, however, their levels could not be linked to vOTU abundance changes. The vOTUs we did detect indicated stress of the *Citrobacter* population, increased bacterial defence systems and potential prophage induction. Given the bacterial data was derived from read profiling, it is possible the increase in *C. koseri* signatures was linked to prophages which were miss-identified as bacterial. The lytic action of the phage cocktail also could have allowed these bacteria to flourish. Although the justusliebigvirus phage bloom observation was the most pronounced change in the system, it is likely that additional phage/host interactions were occurring here without great disturbance. Our gut model and analysis form the basis to investigate changes to bacterial and viral communities in the presence of phage treatment, and has the potential to be utilised with other phage cocktails or formulations. However, our small study would have benefited from more donor samples and replication to better define statistical differences between phage treatment and controls. Furthermore, deeper sequencing and strain-level analyses of metagenome-assembled bacterial genomes would be advantageous in disentangling the intricate interactions and dynamics between the healthy bacteria and phage communities in the gut.

## Conclusions

The commercially available Intesti bacteriophage cocktail preparation may be taken orally to target common bacterial infections of the human gastrointestinal tract, but little is known about how the phages interact with the native gut microbiota. Our simulated gut model was seeded with microbiota consistent with healthy-donor enterosignatures, and our sequencing method enabled us to observe bacterial and viral community changes. Each donor had a unique bacterial composition which altered over time. When comparing phage treated and control samples, *E. coli* had the largest difference in relative abundance and was more closely associated with the controls. This observation suggests lytic action from the phages in the cocktail, such as the justusliebigvirus bloom phage. Changes to the microbiota and impact of phage cocktail treatment were often transient, where we detected minimal changes to typical members of the virome and active prophages in the system. While the full impact of the phage cocktail on the gut microbiota is not known, these initial experiments suggest that the Intesti bacteriophage minimally interrupted the native gut microbiota with the model.

## Acknowledgments

The authors would like to acknowledge Oliver Charity for useful discussions regarding data analysis. The authors acknowledge the use of the QIB Colon Model Facility and thank the manager, Lee Kellingray, for facilitating and advising on this project.

## Funding statements

T.L.B., A.T., R.A. and E.M.A. gratefully acknowledges funding by the Biotechnology and Biological Sciences Research Council (BBSRC) Institute Strategic Programme Gut Microbes and Health BB/R012490/1 and its constituent projects BBS/E/F/000PR10353 and BBS/E/F/000PR10356. E.M.A. is currently funded by the BBSRC Institute Strategic Programme Food Microbiome and Health BB/X011054/1 and its constituent projects BBS/E/QU/230001B and BBS/E/QU/230001D; and the BBSRC Institute Strategic Programme Microbes and Food Safety BB/X011011/1 and its constituent projects BBS/E/QU/230002A, BBS/E/QU/230002B and BBS/E/QU/230002C. R.C. and E.M.A. are funded through the Biotechnology and Biological Sciences Research Council (BBSRC) grant Bacteriophages in Gut Health BB/W015706/1. GS was supported by the BBSRC Core Capability Grant BB/CCG2260/1. J.A.D. was supported by the UKRI Biotechnology and Biological Sciences Research Council (BBSRC) Doctoral Training Partnership BB/M011216/1. CKAE was supported by the Medical Research Council (MRC) as part of the Doctoral Antimicrobial Research Training (DART) MRC iCASE Programme MR/R015937/1. A.T. and R.A were supported by the Quadram Institute Bioscience BBSRC funded Core Capability Grant BB/CCG1860/1.

## Declaration of interest statement

The authors have no competing interests to declare.

## Data availability statement

The raw reads of all shotgun metagenomics experiments are available from the European Nucleotide Archive under Project accession number PRJEB96854. Accession numbers of the individual runs are listed in Supplementary Table 1. A curated database of the assembled viral vOTUs can be accessed on FigShare at the following link: https://doi.org/10.6084/m9.figshare.31016773

## Notes

https://doi.org/10.6084/m9.figshare.31016773

### Competing Interest Statement

The authors have declared no competing interest.

